# LIM Protein Ajuba associates with the RPA complex through direct and cell cycle-dependent interaction with the RPA70 subunit

**DOI:** 10.1101/177840

**Authors:** Sandy Fowler, Pascal Maguin, Sampada Kalan, Diego Loayza

## Abstract

DNA damage response pathways are essential for genome stability and cell survival. Specifically, the ATR kinase is activated by DNA replication stress. An early event in this activation is the recruitment and phosphorylation of RPA, a single stranded DNA binding complex composed of three subunits, RPA70,RPA32 and RPA14. We have previously shown that the LIM protein Ajuba associates with RPA, and that depletion of Ajuba leads to potent activation of ATR. In this study, we show evidence that the Ajuba-RPA interaction occurs through direct protein contact with RPA70, and that their association is cell cycle-regulated and is reduced upon DNA replication stress. We propose a model in which Ajuba negatively regulates the ATR pathway by directly interacting with RPA70, thereby preventing an inappropriate ATR activation. Our results provide a framework to understand the mechanism of regulation of ATR in human cells, which is important to prevent cellular transformation and tumorigenesis.

## Introduction

Cells possess conserved mechanisms that have evolved to sense, respond to and repair specific types of DNA damage (1). These pathways are intimately linked to checkpoint systems and represent highly coordinated and complex responses to extrinsic or intrinsic damage (1). Of those, the ATR pathway responds to DNA replication stress, such as nucleotide imbalance or collapsed forks, or accumulation of single stranded DNA which are events occurring, albeit to a low degree, at each and every S phase of the cell cycle (2) (3). Importantly, the ATR pathway is a tumor suppressor system acting early in tumorigenesis and cell transformation (2). Many aspects of ATR activation have been extensively characterized, involving the accumulation of single stranded DNA binding complex RPA, the phosphorylation on the N-terminus of the RPA32 subunit (4), and further recruitment of the 9-1-1 complex, TOPBP1 and ATR-ATRIP, which are factors necessary for the activation of ATR kinase activity that assemble on the RPA70 subunit (5) (6) (7). The ATR kinase phosphorylates substrates that orchestrate cell cycle pausing and damage repair, such as Chk1 or Claspin. Local recruitment of ATR on RPA includes autophosphorylation at Ser1989 (8), an event that is also required for activation of the pathway. The resulting effects lead to a pause in the cell cycle during S phase, through phosphorylation and activation of Chk1 and Cdc25A, or in some cases apoptosis, through activation of Cdc25C.

RPA plays a crucial role in the ATR activation pathway. RPA is a trimeric complex composed of the three subunits RPA70, RPA32 and RPA14 (9). This complex constitutes an important single stranded DNA binding complex, which binds DNA with high affinity (10) (and ref. therein). DNA binding is mediated by specific domains in RPA70 and RPA32, which adopt a structural pattern called OB-fold (for oligosaccharide/oligonucleotide binding) (11) (12). The trimeric RPA complex possesses six such OB folds, with RPA70 (4), RPA32 (1) and RPA14 (1). The RPA14 subunit is essential for stability of the complex (12). The N-terminal OB fold in RPA70 (termed RPA70N (6)) represents a platform for the assembly of RAD9 and ATRIP-ATR, necessary for the recruitment of TOPBP1, the activator of ATR kinase (6) (13). RPA is essential for DNA replication, by allowing fork progression and lagging strand synthesis, for recombination, by catalyzing strand invasion, and DNA repair, by being involved, among other activities, in ATR activation. In DNA repair, RPA possesses both structural and signaling roles. The complex has a structural role given its ability to bind single stranded DNA, thereby preventing secondary structures incompatible with replication or repair, and a signaling role related to the assembly of the ATR activation complex on RPA70N. We have previously implicated the LIM protein Ajuba as a new player in the ATR pathway (14). The LIM superfamily of proteins, constituted by 60 members in the human proteome, is subdivided into seventeen families, all with predicted LIM domains in various arrangements (15) (16). LIM domains are known protein interaction domains that present distinctive loops defined by interactions between Cys and His residues coordinating a Zn ion (16). Ajuba itself is part of the Zyxin family, which in addition includes TRIP6 and LPP, two components that are involved in telomere protection, through binding of the OB fold protein POT1 (17) (18). The Zyxin family is characterized by the presence of three C-terminal LIM domains (16). We have shown that Ajuba acts as a negative regulator of ATR in unperturbed cells (14). Notably, upon depletion of the protein, cells experience S phase delay along with strong activation of Chk1, a known ATR substrate, and undergo apoptosis. Our current model is that Ajuba exerts an inhibitory activity to full-blown activation of the ATR kinase, and is essential to prevent cell death by apoptosis in unperturbed cells. This inhibitory effect correlates with our observation that Ajuba associates with the RPA complex in the cell (14), and thus associates with a core component of early ATR activation. In this study, we show that the interaction between Ajuba and the RPA complex is direct and occurs through the RPA70 subunit. We found that this association occurs in the nucleus, and that the amounts of Ajuba along with the degree of co-localization between Ajuba and RPA are increased in S phase, suggesting cell cycle-dependent regulation of the localization of Ajuba, and perhaps of its interaction with RPA as well. We show that hydroxyurea, known to induce DNA replication stress and activation of ATR, leads to dissociation of Ajuba from RPA and to efficient export of Ajuba from the nucleus. Based on our findings, we propose a model for Ajuba in preventing inappropriate ATR induction during S phase.

## Results

### Ajuba associates with the RPA complex and dissociates upon replication stress

In order to determine whether Ajuba exhibits association with the whole RPA complex, or to a particular subunit of RPA, co-immunoprecipitations were performed with HTC75 cell extracts using antibodies to each subunit separately. As seen in Figure 1A, Ajuba is able to immunoprecipitate RPA70 and RPA32. In addition, antibodies to each of the three RPA subunits are able to pull down Ajuba (Fig.1B). This suggests that Ajuba associates in cells with the whole RPA complex, and not with a particular subunit separately. In each case, the estimate of the pull down efficiency is between 2 and 5% of total RPA in association with Ajuba (see Materials and Methods). Therefore, a small pool of RPA is involved in the association. In turn, a small fraction of Ajuba is in association with the RPA complex. Therefore, the interaction is robust but involves a small fraction of either partner.

**Figure 1.**
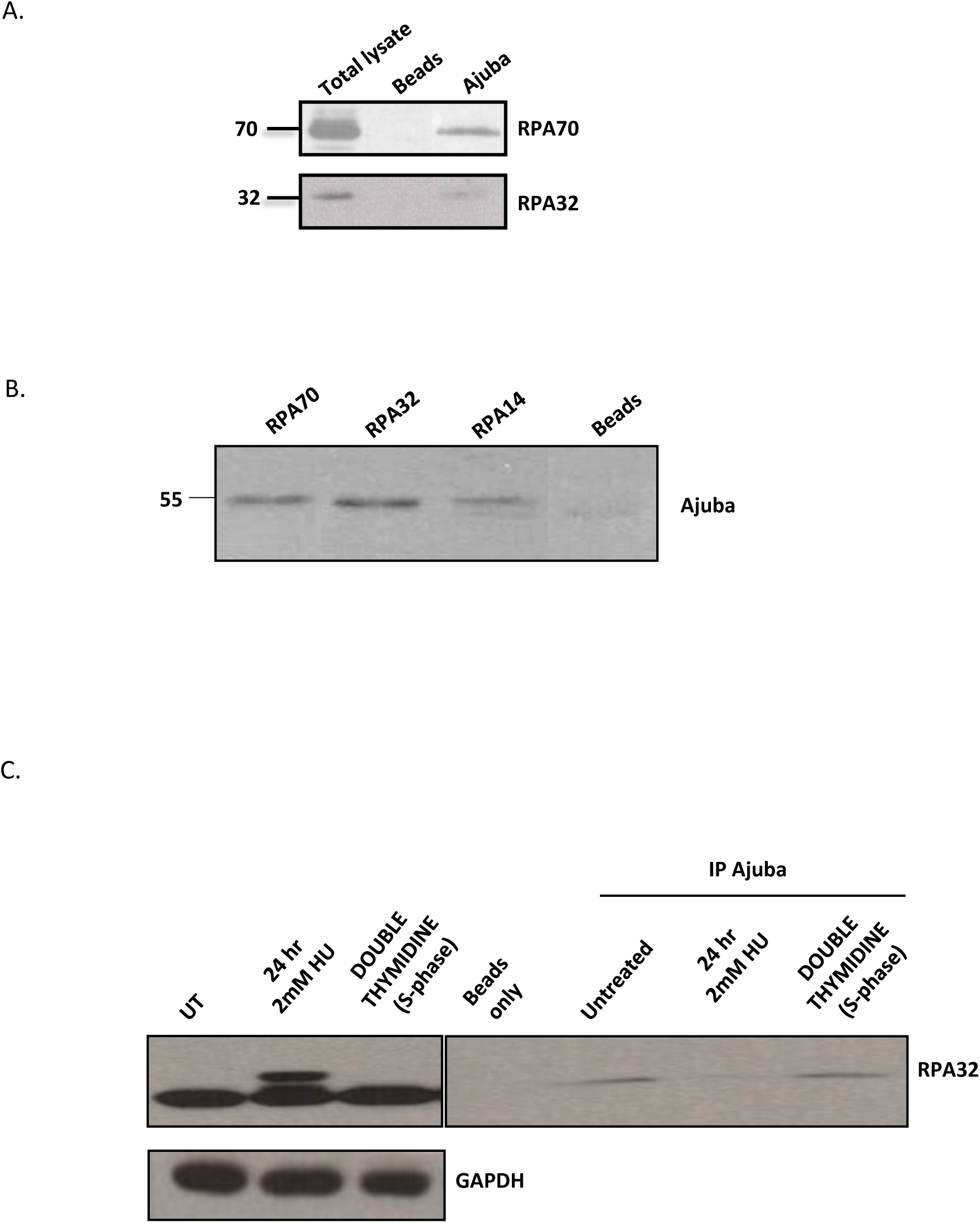
Ajuba is in association with the RPA complex, and dissociates upon hydroxyurea treatment. A) IP-Western on immunoprecipitations with Ajuba serum from HTC75 total extracts. The blots were probed with the indicated RPA antibodies. B) IP-Western using the indicated RPA antibodies for immunoprecitations, and blotted for Ajuba. C) IP-Western with anti-Ajuba immunoprecipitations, and probed for RPA32. Total extracts shown on the left panel. Cells were untreated (control), treated with 2mM HU for 24 hr, or synchronized by double Thymidine block and released for 2 hr. See also figures S1, S2 and S3 for data with IMR90 cells.

Our previous studies showed that depletion of Ajuba led to an apparent activation of ATR in otherwise unperturbed cells (14). We asked whether the enforcement of the ATR response through DNA replication stress induced by hydroxyurea treatment had an effect on the association. We found that treating cells with 2mM HU led to a significant reduction in the association of Ajuba and RPA, as observed by co-immunoprecipitation with RPA32 (Fig.1C, right panel). The activation of the DNA damage response is evident from the induction of phosphorylation of RPA32 (1C, left panel). The same effect of hydroxyurea of reducing the pull down of RPA by Ajuba is seen for RPA70 (Fig.S1), arguing for a dissociation of Ajuba with the whole RPA complex under condition of replication stress. In addition, the overall amounts of Ajuba are unchanged during hydroxyurea treatment (Fig.S4), showing that the decrease in the amounts of Ajuba is not due to a reduction in protein stability. Since hydroxyurea treatment influences the cell cycle profile in culture, by enriching for cells in S phase (Fig. S3), we asked whether the dissociation of Ajuba and RPA was a result of cell cycle stage. We therefore synchronized cells in G1 through a double thymidine block, released the cells in S phase and probed for possible Ajuba-RPA dissociation (Fig.1C, right panel). We found no perceptible effect on the amount of Ajuba associated with RPA in mostly S phase cells, and concluded that the dissociation seen upon hydroxyurea treatment was not merely due to delay in S phase, and therefore was the consequence of DNA replication stress. Finally, the association of Ajuba with RPA and the effect of hydroxyurea are not a specific cell-line or tumor phenotype, since the same observations were done using IMR90 cells, a normal diploid human fibroblast cell line (Figs. S1, S2, S3).

### Ajuba nuclear localization is cell cycle regulated, and is significantly increased in S phase

Whereas the RPA complex is localized in the nucleus, Ajuba possesses a complex intracellular trafficking: as other members of the Zyxin LIM family, it is mostly present in the cytoplasm, where it is involved in cell-cell and cell-matrix adhesion, and is actively shuttled in and out of the nucleus by virtue of a discernible nuclear export sequence (NES). Thus, a small pool of Ajuba (about 10%, S.K. and D.L., unpublished) is present at steady state in the nucleus. Given the dynamic trafficking of Ajuba, and our observation that it controls ATR activation in S phase, we probed whether the localization of Ajuba was dependent on the cell cycle. In unsynchronized cells, about 33% of the cells showed a clear nuclear staining pattern for Ajuba as shown in Fig. S5A, in HTC75 and IMR90 cells. In those, the staining pattern for Ajuba exhibited clearly distinguishable foci, overlayed on an overall diffuse nuclear staining pattern. In cells synchronized and released in S phase, we found a clear increase in the fraction of cells containing nuclear Ajuba (Fig.2): the percentage increased to 68.9% at the 4-hr time point, corresponding to mid-S phase, and was followed by a decrease to 46.8% at the 6hr time point (Fig.2). The 0 hr time point (G1 cells) showed no significant difference with unsynchronized cells, arguing that the 33% of cells with nuclear Ajuba correspond mostly to nuclear Ajuba in G1. Therefore, we conclude that a fraction of Ajuba is present in the nucleus throughout the cell cycle, but that a significant increase in nuclear accumulation occurs during S phase. This increase could occur through a combination of effects on the flow of nuclear import, and the efficiency of export through the NES in the molecule. We do believe that nuclear export is an important step because of the observation that Leptomycin B, an inhibitor of CRM1-mediated export, leads to a significant increase in Ajuba’s nuclear localization (up to 59.3%, see Fig. S5B and S5C).

**Figure 2.**
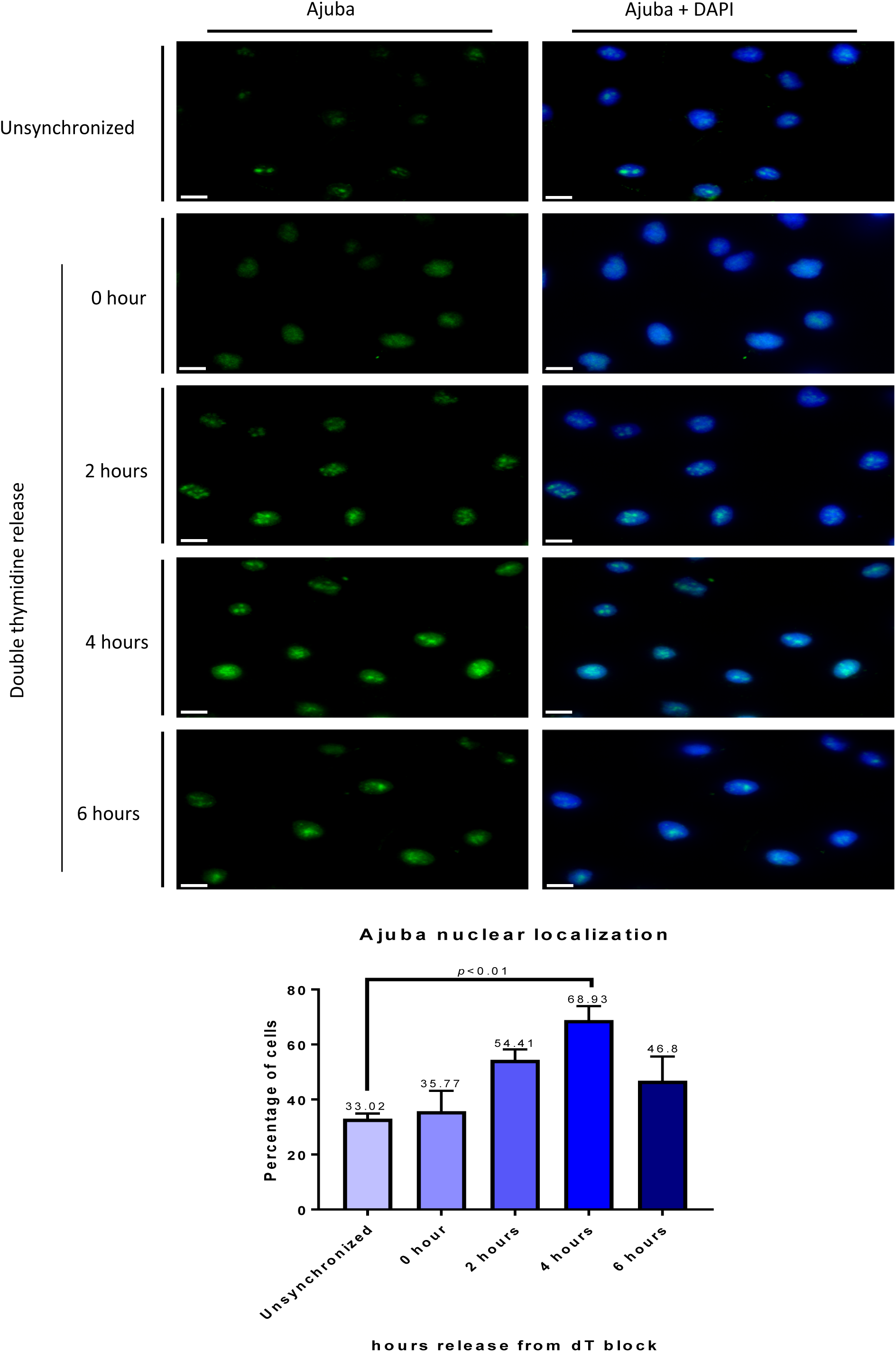
Ajuba nuclear localization is increased during S phase. HTC75 cells were synchronized to G1/S border with double thymidine block and released into S phase, with cells processed for IF at the indicated time points. (Top) Immunofluorescence of Ajuba in unsynchronized and synchronized cells. (Bottom) Quantitation of cells positive for nuclear Ajuba at each time point (n=100).

### Ajuba partially co-localizes with RPA in the nucleus, with co-localization increased in S phase

To address whether Ajuba-RPA interaction takes place in the nucleus, we performed co-immunofluoresence for both proteins in HTC75 and IMR90 cells. We found a significant degree of co-localization in the nucleus, in unsynchronized cells (Fig.S5A). The co-localization was observed specifically with the Ajuba signal that was found in foci, and only in a subpopulation of cells. Given our result that Ajuba appeared enriched in S phase nuclei, we asked whether the co-localization between Ajuba and RPA occurred primarily in S phase. We therefore repeated the experiment with cells released in S phase, and found that the percentage of nuclei exhibiting more than 3 nuclear foci for both Ajuba and RPA70 greatly increased up to mid S phase (4hr time point), from 17.6% to 61.9%, then decreased to 27.4% (6hr time point) (Fig.3). Again, cells in G1 (0hr) did exhibit some level of co-localization (17%), but to a lesser degree compared to unsynchronized cells (17.6% versus 27.5%). Thus, the amounts of Ajuba and degree of association with the RPA complex are not exclusive to S phase, but appear to be highly enriched at an interval representing early to mid-S phase. In each nucleus, the degree of co-localization in foci was not complete, indicating that some sites containing Ajuba remain to be identified. We looked at the possibility of co-localization between and Ajuba and PCNA, essential for lagging strand synthesis. We did find significant co-localization between Ajuba and PCNA as well (Fig.S6). It is therefore possible that some of the nuclear Ajuba foci represent sites of active DNA replication.

**Figure 3.**
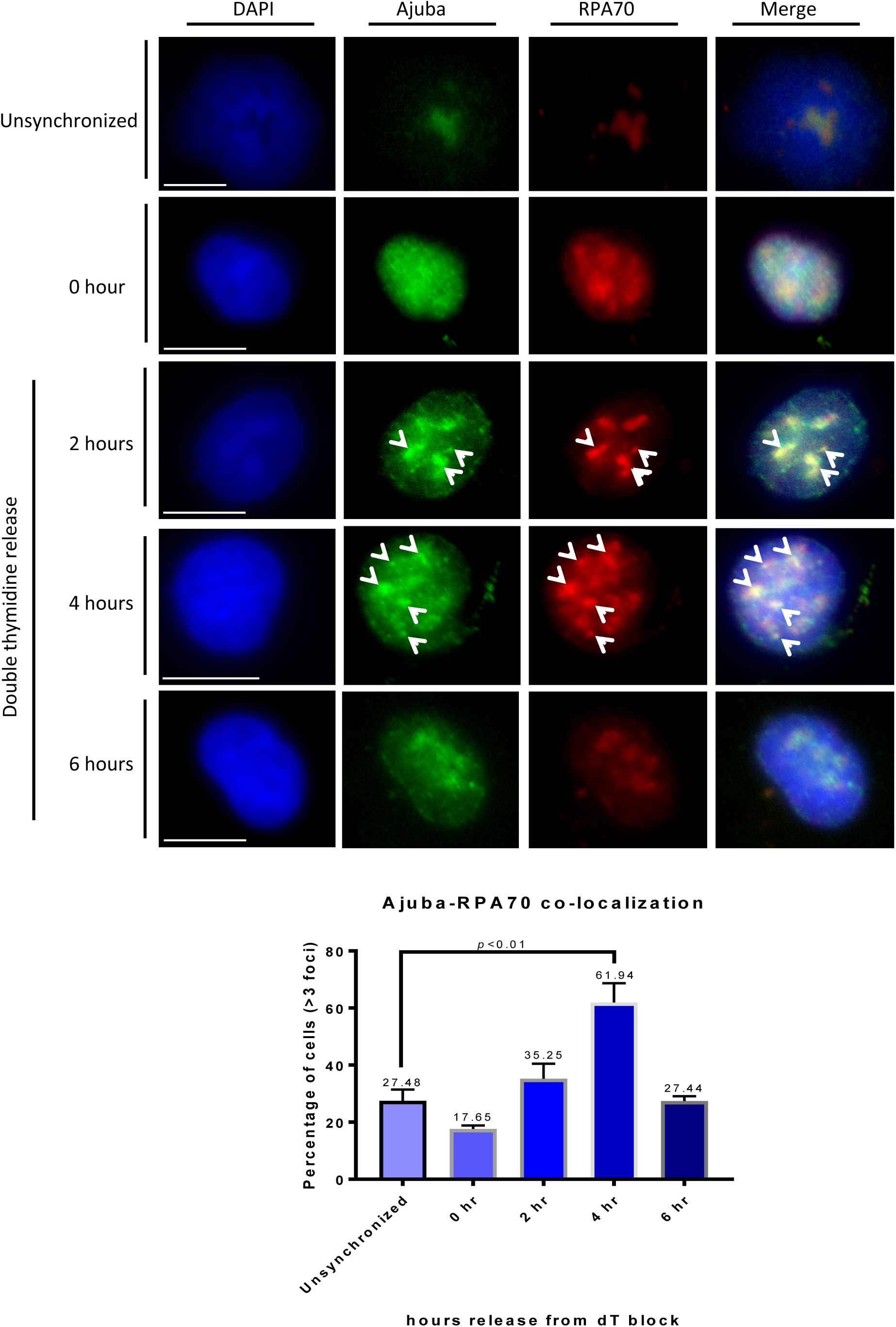
Increase of Ajuba-RPA70 co-localization in the nucleus during S phase. HTC75 cells were synchronized to G1/S border with double thymidine block and released into S phase with cells processed for IF at the indicated time points. (Top) Co-immunofluorescence of Ajuba and RPA70 in unsynchronized and synchronized cells. Arrowheads point to sites of co-localization (Bottom) Quantitation of cells exhibiting > 3 foci of Ajuba-RPA70 co-localization in the nucleus at each time point (n=100).

### Ajuba interacts with the RPA70 subunit directly

The association between Ajuba and the RPA complex could be indirect and mediated by additional factors, or direct through protein-protein interaction between Ajuba and one of the subunits. We investigated whether a direct protein contact could be detected between Ajuba and one of the RPA subunits. To this end, the *in vitro* translation system using rabbit reticulocyte lysates was employed to address the nature of interaction between Ajuba and RPA. Each of the RPA subunit was cloned into pCMVTnT vector and Ajuba was cloned into a N-terminal His-tagged vector (pcDNA 3.1/His B). Ajuba was co-translated with each of the RPA subunits separately and the reaction mixtures were subjected to His-tag pulldown using nickel beads. As shown in Figure 4A, Ajuba was found to pull down with the large subunit of the RPA complex, RPA70. Quantitative analysis of pull down efficiencies showed that 61.9% of the *in vitro* translated RPA70 was precipitated by HIS-Ajuba, with RPA32 yielding only background (13%) (Fig.4B). As a control, we probed whether Ajuba could associate with POT1, the OB fold telomeric single strand binding protein. There was no detectable binding between the two (Fig. S8). Our data indicate that there is a direct and specific contact between Ajuba and the RPA complex, through the RPA70 subunit.

**Figure 4.**
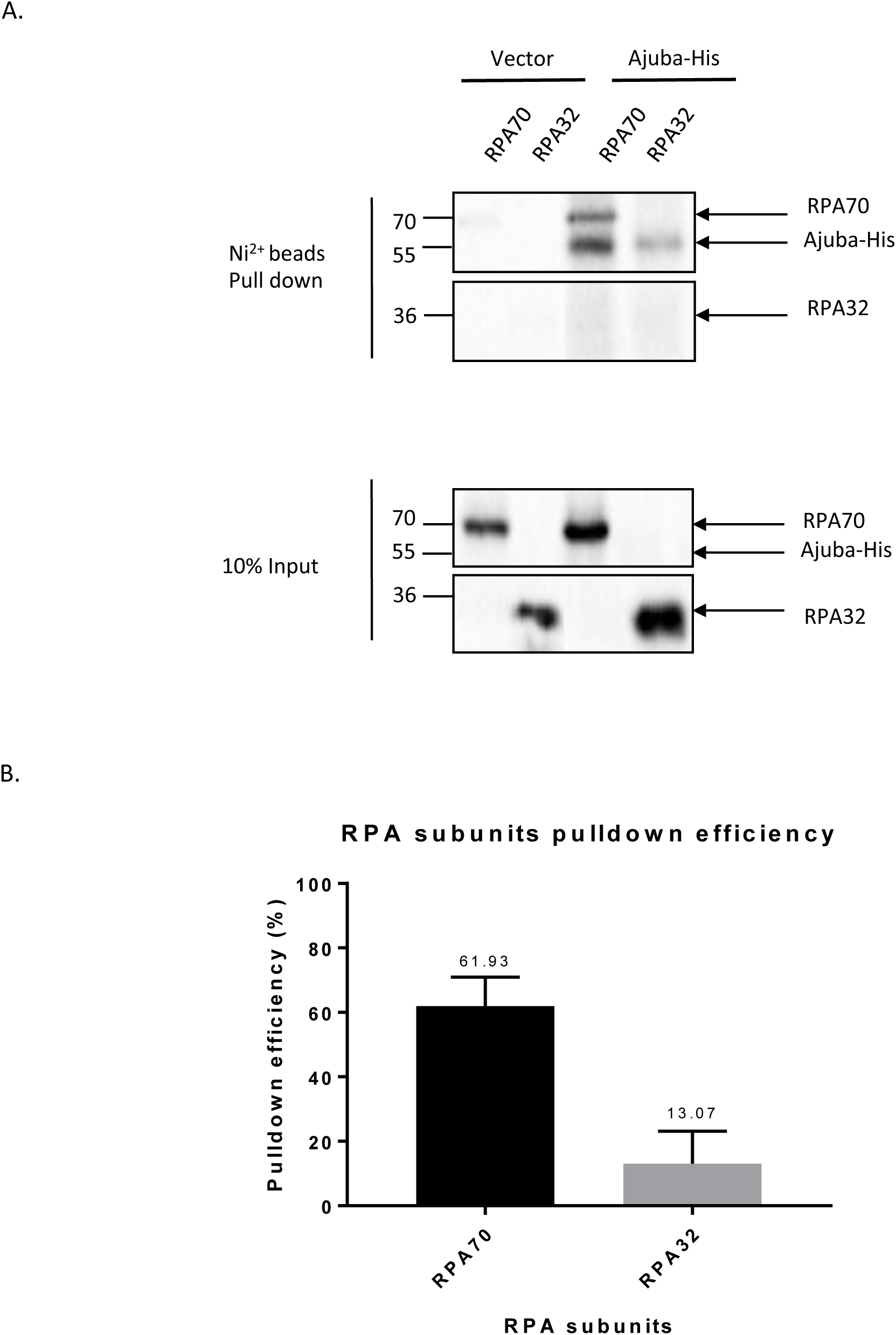
Ajuba can bind directly to the RPA70 subunit in vitro. *In vitro* co-translation of His-tagged Ajuba (Ajuba-His) and RPA subunits: RPA70, RPA32, and RPA14 (not shown). (Top) Autoradiography of affinity pulldown with Ajuba-His and co-translated RPA subunit. (Bottom) Pull down efficiency (%) of each RPA subunit by Ajuba-His. Pull down efficiencies were determined by the quantification of pull down signal in autoradiography and normalized to input signal and the number of methionine of each construct. See Materials and Methods for further details.

### Direct contact between RPA70 and Ajuba is mediated by OB fold A of RPA70 and the central region of Ajuba

The Ajuba-RPA70 interaction prompted us to further define the region on both proteins that mediate this interaction. In order to map the region of interaction, truncation alleles of RPA70 were generated by PCR and cloned into pCMVTnT vector. The N-terminus of RPA70, OB fold F, has been extensively studied for its function as an “interaction hub” with proteins involved in DNA damage and checkpoint response (6). In the central region of RPA70, OB folds A and B have been shown to associate with proteins involved in DNA replication and repair. At the C-terminus, OB fold C was shown to be involved in subunit interaction (12). Truncations were constructed based on previous sequence and structural analysis, hence separating RPA70 into OB fold F and OB folds A-B-C. The alleles were co-translated with His-tagged full-length Ajuba and the reaction mixtures were subjected to His-tag pull downs.

Figures 5A and 6A show the pull downs between full-length Ajuba and different RPA70 truncations as shown, and the quantification of the pull down efficiency of each allele. OB folds A-B-C was found to pull down with full-length Ajuba with a yield very similar to full-length RPA70 (Fig. 5A). When half of OB fold A was truncated, the pull down with full-length Ajuba was decreased to 58.7%, with full-length RPA70 being set at 100% (Fig. 5B, 6A). OB fold F, as well as an allele containing only OB folds B and C, showed minimal binding activity with full-length Ajuba (Fig. 6A, S7). Taken together, Ajuba physically contacts OB ABC of RPA70 and not OB F *in vitro*. In particular, OB fold A seemed to be the most important region for the interaction.

**Figure 6.**
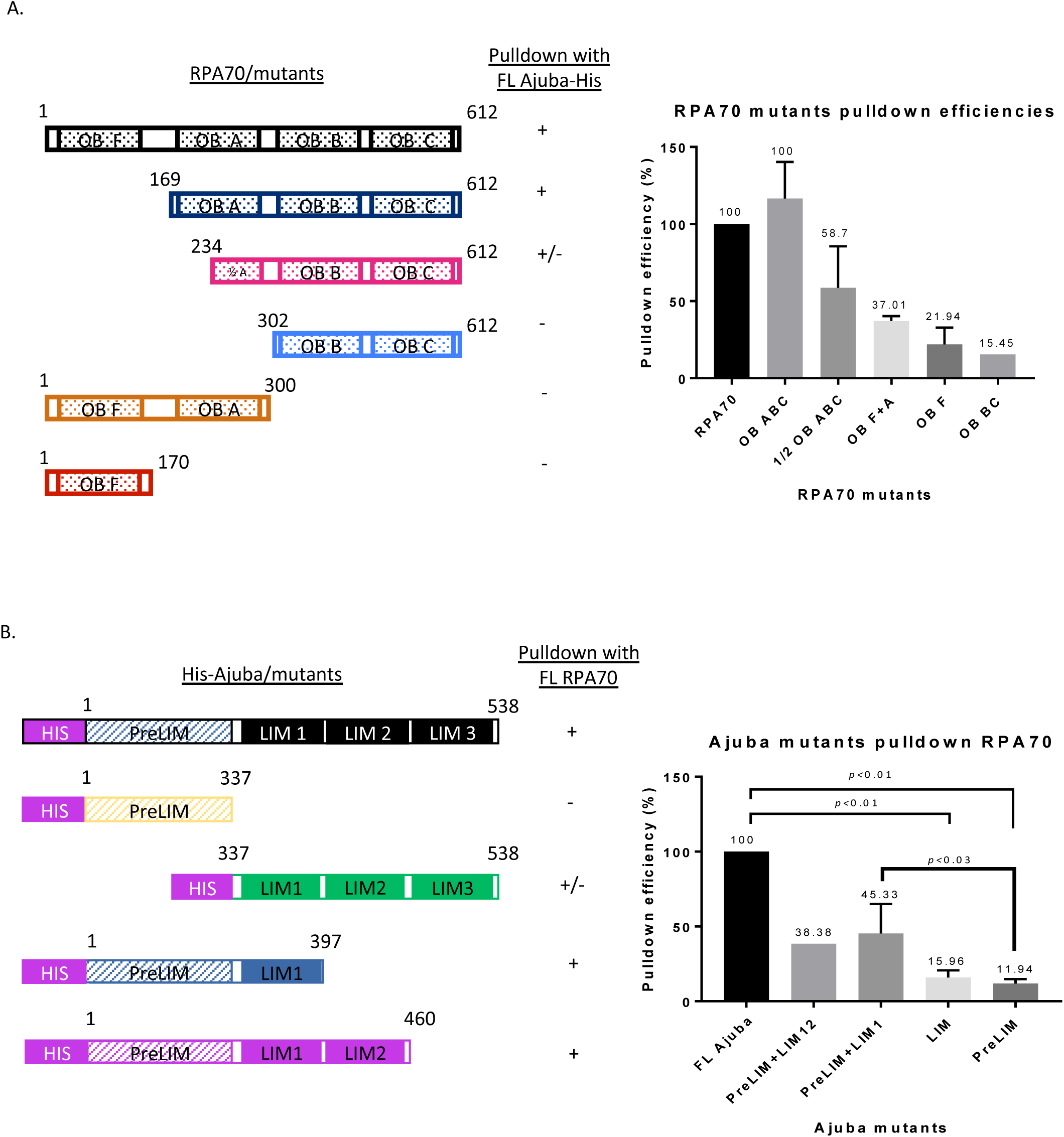
RPA70 and His-tagged Ajuba truncation allele constructs and pulldown efficiencies *in vitro.* A. (Left) Full-length RPA70 and truncation constructs. (Right) Pull down efficiencies of each RPA70 by full-length Ajuba-His. B. (Left) Full-length Ajuba-His and His-tagged truncation constructs. (Right) Pull down efficiencies of full-length RPA70 by each His-tagged Ajuba truncation allele.

**Figure 5.**
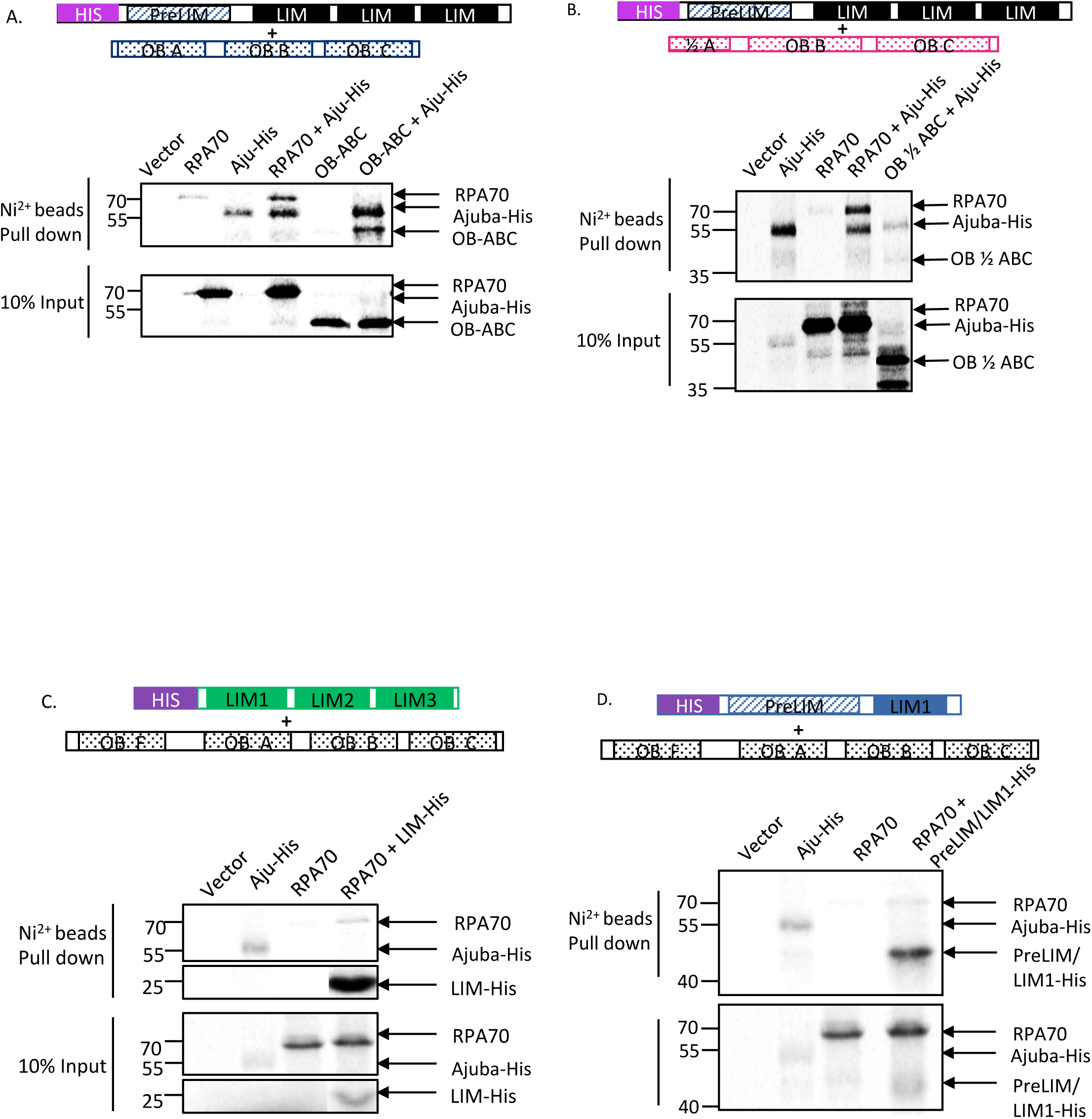
RPA70 OB fold A and Ajuba LIM domain 1 are important, but not sufficient, for the Ajuba-RPA70 interaction *in vitro.* Autoradiography of His tag pull downs with nickel beads in Ajuba/RPA70 mutants co-translations. A. Deletion of OB fold F does not impact pull down with full-length Ajuba-His. B. Truncation of half of OB fold A reduced pull down with full-length Ajuba-His. C. LIM 1-2-3 pulls down full-length RPA70. D. Addition of LIM domain 1 to the preLIM region enhanced pull down of full-length RPA70.

We attempted to test whether OB fold A was sufficient for the interaction by testing an allele fusing OB fold A with OB fold F. Since OB F did not exhibit interaction with Ajuba, any binding activity would be ascribed to OB fold A. Surprisingly, OB F+A did not exhibit any major increase in binding with Ajuba compared to OB F alone (Fig 6A). Collectively, our data suggests that OB A is necessary but not sufficient for the interaction with Ajuba, and possibly other domains in RPA70 are required to reconstitute full binding activity.

We also sought to identify interaction domains in Ajuba. A previous study has described the preLIM region by itself localized in the cytoplasm, and LIM domains bearing a determinant for nuclear localization (19) (20). This prompted construction of alleles containing the preLIM region and LIM domains, and probed for RPA70 interaction. Each allele was co-translated with full-length RPA70 and subjected to binding assays with nickel beads. Figure 5C and 5D show that both Ajuba alleles exhibit very low binding activity.

We hypothesized that the actual binding domain in Ajuba was severed in our initial constructs, and might lie between the pre-LIM and LIM regions. We then constructed additional alleles with different end points, specifically harboring the preLIM region with adjacent LIM domains (LIM 1 and 2) in order to preserve the central region. As shown in Fig. 5D and 6A, the pull down signal of full-length RPA70 increased when LIM domain 1 or LIM domains 1 and 2 are added to the preLIM region. Quantitation of RPA70 pulldown efficiency by Ajuba mutants (Figure 6B) shows a significant decrease in binding activity in all mutants compared to full-length Ajuba, with pre-LIM+LIM1 being the most efficient at 45.3% of full-length (Fig.6B), once again arguing that no single domain could account for the full binding activity of the protein. In addition, prelim-LIM1 and preLIM-LIM1-LIM2 constructs exhibited higher pull down efficiencies than either preLIM region or LIM domains alone. This suggests that the central region of Ajuba participates in this physical contact with RPA70, along with other possible motifs. Our model then is that the central portion in Ajuba, between residues 337 and 397, mediates a direct interaction with OB fold A of RPA70, between residues 169 and 300. Our model for this interaction is shown in Fig. S9.

### Hydroxyurea leads to significantly diminished nuclear accumulation of Ajuba

The decreased association of Ajuba with RPA in cells experiencing DNA replication stress, combined with the enrichment of Ajuba in S phase nuclei prompted us to ask whether hydroxyurea could influence the intracellular localization of Ajuba: possibly, the dissociation of Ajuba from RPA could lead to active export in conditions of ATR activation. Cells were treated with 2mM hydroxyurea for 24hr and the localization of Ajuba was assessed. We found that the nuclear accumulation of Ajuba was greatly reduced, from 38% in controls to 16.7% in cells treated with 2mM HU (Fig.7). The concomitant change in the staining pattern of RPA 70 in 2mM HU confirmed the induction of DNA replication stress in nuclei with lowered Ajuba accumulation. Altogether, our data suggest that DNA replication stress leads to dissociation of Ajuba from RPA and active export from the nucleus.

**Figure 7.**
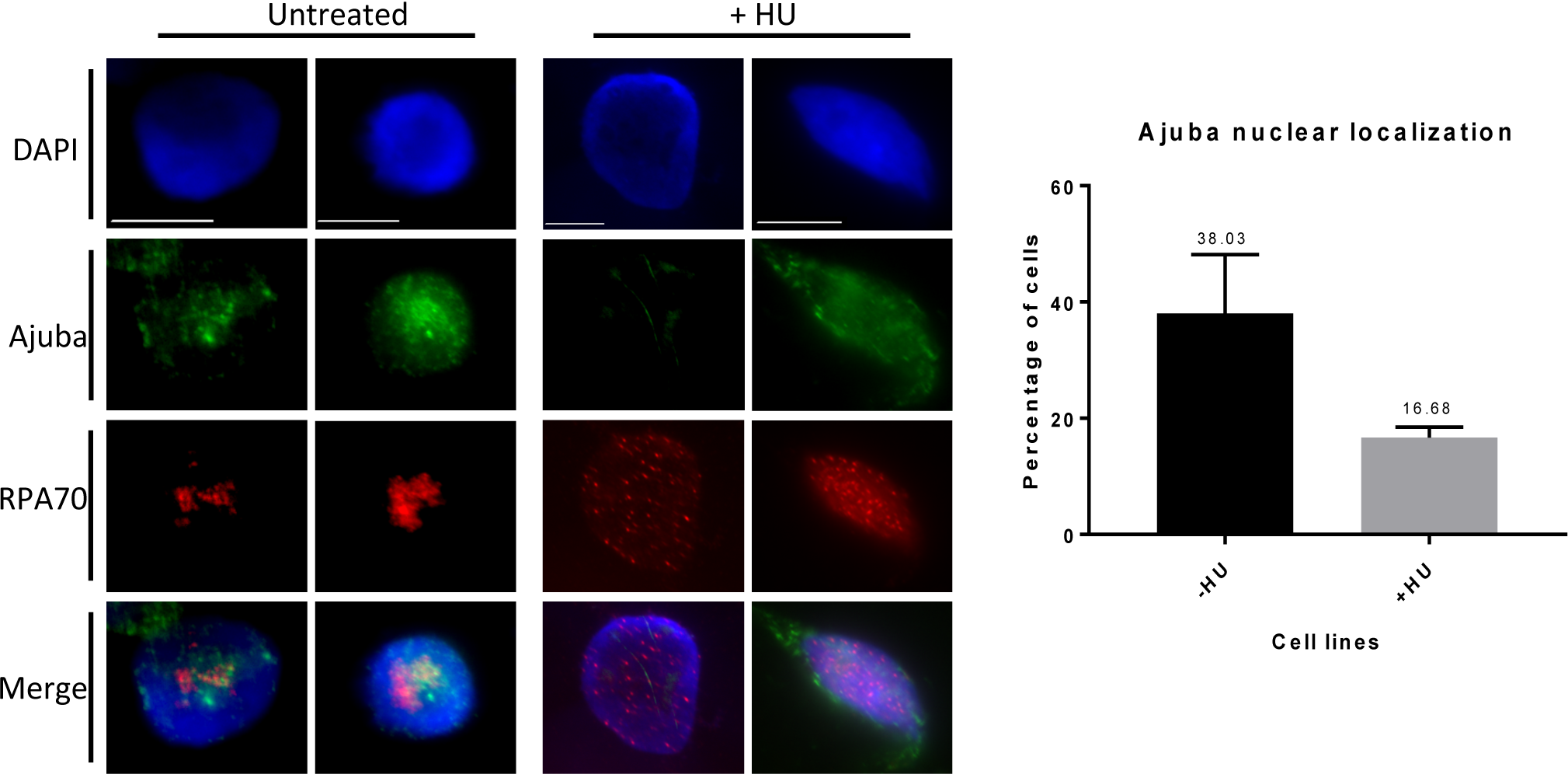
Ajuba nuclear localization is reduced upon HU treatment. (Left) Co-immunofluorescence of Ajuba (FITC) and RPA70 (TRITC) in untreated and HU treated HTC75 cells. (Right) Quantification of cells exhibiting strong nuclear localization with and without HU treatment (n=100, three independent experiments).

## Discussion

A number of studies have established a well-understood molecular pathway for ATR activation: stabilization of RPA on single stranded DNA, present in excess to normal amounts produced by the replication forks in S phase, leads to the assembly of a recruitment complex on the N-terminal OB fold of RPA70 (called OB-F or RPA70N) (6) (7). The RPA70 subunit, therefore, represents a platform onto which RAD9, part of the activation 9-1-1 complex (RAD9-RAD1-HUS1), TopBP1 and ATR-ATRIP are recruited, leading to autophosphorylation of ATR and induction of the kinase (8). While the assembly of the ATR activation complex can occur *in vitro* in a purified system, little is know about how ATR activation is prevented from occurring under non-inducing conditions. We have proposed, based on an earlier study, that Ajuba plays an important role in this context (14). Given the observations that depletion of Ajuba leads to the induction of ATR in absence of any damage or causative agent, we have proposed that Ajuba acts as an inhibitor of ATR induction. We have also discovered that Ajuba associates with RPA, placing it at the core of ATR activation. Here, we show that the association between Ajuba and RPA is direct, through a contact with RPA70, and that, in cells, activation of ATR by hydroxyurea leads to dissociation of Ajuba from RPA accompanied by a strong reduction in nuclear accumulation. Our view is that the ATR activation complex is poised for assembly on the RPA70 recruitment platform, and that Ajuba must be dissociated from RPA for activation to occur. Whether Ajuba acts specifically in S phase remains to be determined, but our data suggest that its activity is especially important for S phase progression, given that depletion of the protein eventually leads to strong apoptotic signals, with the remaining cells being mostly delayed in S phase.

Ajuba has been described in a different setting, in maintaining the structure of desmosomes at cell adhesion sites (21). Indeed, the majority of the protein is extra-nuclear, with only a small fraction (10% or less) present in the nucleus, and this in a small subset of the cells (SK and DL, unpublished, this study). Ajuba and other members of the Zyxin subfamily (Zyxin, TRIP6, LPP) are known to undergo a dynamic intracellular life cycle, with active shuttling in and out of the nucleus (22) (see our model, Fig.8). Export from the nucleus is an important part of this life cycle, because interfering with nuclear export leads to significant protein accumulation in the nucleus (23) (24). Based on our results, the pool of Ajuba present in the nucleus could be maintained there through direct binding to RPA70, in the context of ATR inhibition (Fig.8). Whether the intra- and extra-nuclear roles of Ajuba are linked and pertain to some kind of subcellular signaling remains to be determined, and separation-of-function alleles would be informative in this regard. Our work represents a first step in the design of such alleles. Once DNA damage has occurred, Ajuba could dissociate from RPA70 leading to active NES-dependent export out of the nucleus, possibly by “unmasking” of this motif following dissociation.

**Figure 8.**
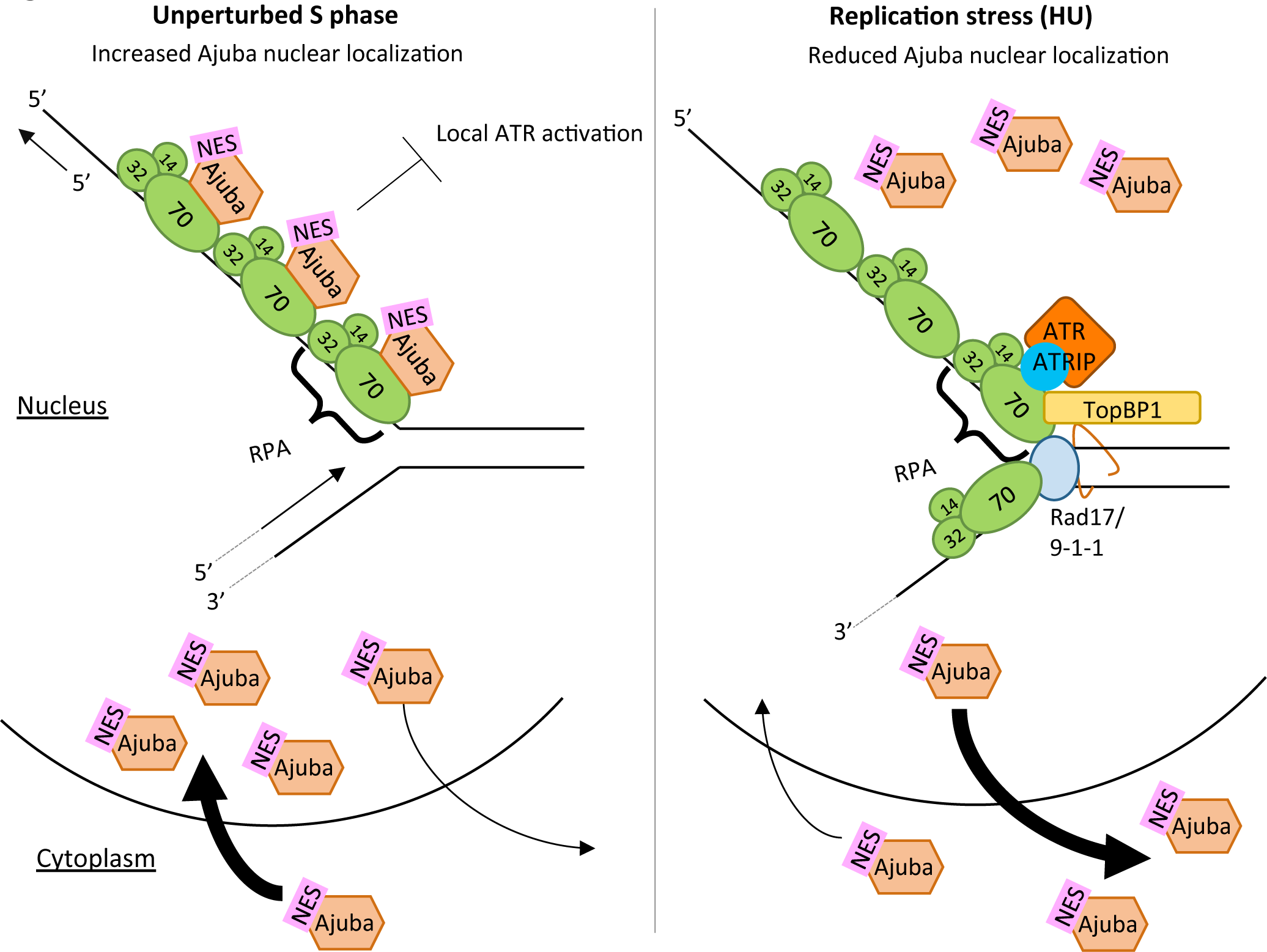
Model for Ajuba’s function during unperturbed S phase and replication stress (induced by HU treatment). (Left) During unperturbed S phase, Ajuba nuclear localization increases, and Ajuba associates with the RPA heterotrimer in the nucleus through direct interaction with RPA70-OB fold A. This interaction inhibits local ATR activation at the replication site. (Right) During replication stress (induced by HU treatment), Ajuba dissociates from RPA and allows for the assembly of the ATR-inducing complex. Ajuba is shuttled out of the nucleus via its NES (nuclear export signal). ATR activation takes place at the specific site of damage, but not at other sites in the genome where Ajuba continues to exert its repressive activity.

Our model put forth previously (14) proposed that Ajuba prevents activation of ATR under non-inducing conditions, activation which occurs upon Ajuba depletion by siRNA, generating by itself conditions of induction in absence of DNA damage. The negative regulation exerted by Ajuba on the ATR pathway could occur by steric hindrance of the assembly of the ATR activating complex on RPA70. Dissociation of Ajuba or its depletion could free up the recruiting surface for the 9-1-1 complex, TOPBP1 or ATR-ATRIP and allow for spontaneous activation downstream. More work would be required to test this model, by asking whether binding of Ajuba to RPA70 is mutually exclusive with components of the ATR activation complex *in vitro*. Alternatively, Ajuba could reduce the affinity of RPA for the DNA, thereby leading to a decrease in RPA-ssDNA complexes and a higher threshold of ATR activation. In this case, analyzing the affinity of the RPA complex for DNA in the presence of Ajuba would be informative. This leaves an important question: how can ATR be activated in the first place? We propose that activation conditions (such as hydroxyurea) lead to dissociation of Ajuba. What leads to this dissociation, or reduction of binding affinity of Ajuba for RPA70, remains to be elucidated. Phosphorylation of RPA32 could be an important event in this context, by preventing Ajuba from reassociating with the complex, or by promoting Ajuba dissociation from the complex, thereby reducing the occupancy of the activating platform on RPA70. Another possibility could be that DNA damage induces a posttranslational modification of Ajuba itself, resulting in a decrease in affinity for RPA. These are testable models for future studies. Finally, our model proposes that Ajuba is involved in local activation of the kinase, and is important to prevent a global response: it could explain activation of ATR at a single replication fork, or at a specific site of DNA damage in the genome, and how the response is elicited locally, while kept at bay globally elsewhere in the genome. We propose that this type of regulation is essential to cell viability in order to impart the right dosage of a response essential to genome integrity, but which, when hyper-activated, would lead to cell death by apoptosis. Ajuba would then allow a “fine tuning” of the ATR response, by preventing the ATR pathway to be activated at places other than the actual site of damage and keeping the response locally constrained. It would be interesting to address whether Ajuba is implicated in other RPA-dependent responses, such as homology-directed repair, or interstrand crosslink repair (25).

## Materials and Methods

### Recombinant protein plasmid construction

For *in vitro* translations Ajuba and Trip6 was amplified using PCR with primers, Ajuba-HIS-5’ and Ajuba-HIS-3’, digested with EcoRI and Xhol, and cloned into pcDNA 3.1/His-B. RPA subunits were amplified with PCR using primers with EcoRI digestion sites and cloned into pCMVTnT vector (Promega). RPA70-5’(5’-GTA TAT GAA TTC ATG GTC GGC CAA CTG AGC GAG-3’), RPA70-3’ (5’-GTA TAT GAA TTC TCA CAT CAA TGC ACT TCT-3’), RPA32-5’ (5’-GTA TAT GAA TTC ATG TGG AAC AGT GGA TTC GAA-3’), RPA32-3’ (5’-GTA TAT GAA TTC TTA TTC TGC ATC TGT GGA-3’), RPA14-5’ (5’-GTA TAT GAA TTC ATG GTG GAC ATG ATG GAC TTG-3). RPA14-3’ (5’-GTA TAT GAA TTC TCA ATC ATG TTG CAC AAT-3’). POT1 was amplified with PCR with primers containing Xhol digestion sequence into pCMVTnT vector. POT1-5’ (5’-GTA TCC TCG AGA TGT CTT TGG TTC CAG CAA C-3’), POT1-3’ (5’-GTA TCC TCG AGT TAG ATT ACA TCT TC TGCA AC-3’).

Ajuba truncation mutants were produced using PCR with primers containing EcoRI and Xhol digestion sites into pcDNA3.1-B/NT-His vector (Invitrogen). LIM-HIS, aa 337-538: LIM-HIS-5’ (5’-CAC ACG AAT TCT GGC ACC TGT ATC AAG TGC AAC-3’), Ajuba-HIS-3’. PreLIM-HIS, aa 1-337: Ajuba-HIS-5’, PreLIM-HIS-3’ (5’-GTA CAC CTC GAG TCA GCC GAA GTA GTC CTC CCT GGC-3’). PreLIM/LIM1-HIS, aa 1-397: Ajuba-HIS-5’, PreLIM/LIM1-HIS-3’ (5’-GTA CAC CTC GAG TCA CTG AAA CCC TGA AAA CAG-3’). PreLIM/LIM12-HIS, aa 1-460: Ajuba-HIS-5’, PreLIM/LIM1+2-HIS-3’ (5’-GTA CAC CTC GAG TCA AGC ATA ATT TTT GTG GTA-3’).

RPA70 truncation mutants were produced using PCR with primers containing EcoRI digestion sequences into pCMVTnT vector. RPA70-OBABC, aa 169-612: OBABC-5’ (5’-CAC AAC GAA TTC ATG GGT CCC AGC CTG TCA CAC-3’), RPA70-3’. RPA70-OBF, aa 1-169: RPA70-5’, OBF-3’ (5’-CAC AAC GAA TTC TCA TGC AGC TTT TCC AAA TGT CTT-3’). RPA70-OBB+C, aa 302-612: OBBC-5’ (5’-CAC AAC GAA TTC ATG GAT TTC ACG GGG ATT GAT GAC-3’), RPA70-3’. RPA70-OBF+A, aa 1-169: RPA70-5’, OBF+A-3’ (5’-CAC AAC GAA TTC TCA GAA ATC AAA CTG AAC CGT-3’). RPA70-OB½ABC, aa 234-612: OB½ABC-5’ (5’-CAC AAC GAA TTC ATG CGA GCT ACA GCT TTC AAT-3’), RPA70-3’. Clones were confirmed by sequencing.

### Cell culture

HTC75 and IMR90 (human diploid lung fibroblasts) (ATCC^®^ CCL-186^tm^) cell lines were employed. HTC75 is a derivative of HT1080 (human fibrosarcoma). IMR90 was used at population doubling 30. HTC75 cells were cultured in DMEM (Cellgro) with 10% BCS (HyClone), 1% penicillin and streptomycin (Cellgro) and 1% L-glutamine (Gibco). IMR90 cells were cultured in DMEM with 20% FBS (ClonTech), and 1% penicillin and streptomycin. Both cell lines were grown in cell culture incubator at 37°C with 5% CO_2_.

### Hydroxyurea treatment and double thymidine block

Hydroxyurea (Thermo Scientific) was added to the medium to a final concentration of 2mM. Cells were collected after 24 hours of treatment. Double thymidine block was performed by adding 2mM thymidine (final concentration) to the media. Cells were synchronized for 18 hours, rinsed with PBS, and fresh media was added to release for 10 hours. Thymidine (2mM final concentration) was added the second time for 18 hours. Cells were rinsed with PBS and fresh media was added to release cells from G1/S border for hours indicated.

### Co-Immunoprecipitation

Cell extracts preparation and protocol were carried out as described in (26).

### Co-immunofluorescence

HTC75 and IMR90 cells were grown on glass coverslips and were washed twice with PBS at room temperature. Nucleoplasmic proteins were extracted with Triton X-100 buffer at room temperature for 10 minutes. Cells were rinsed with PBS twice and fixed in 3% paraformaldehyde/2% sucrose in PBS for 10 minutes at room temperature. Cells were washed with Triton X-100 buffer for 10 minutes at room temperature and washed with PBS twice (5 minutes each). Blocked with PBG for 30 minutes at RT. Added primary antibody diluted in PBG (Ajuba 1:2500, RPA70 1:2500) and stored at 4C overnight. Washed with PBG three times and incubated with fluorescent secondary antibody diluted in PBG for 45 minutes at RT (Anti-rabbit 1:1000, Anti-mouse 1:1000). Washed with PBG twice and incubated the coverslip with DAPI in PBG at 100 ng/ml. Coverslips were mounted on microscope slides with embedding media and sealed with nail polish. Antibodies used: Ajuba (rabbit, Abcam), RPA70 (mouse, Abcam), PCNA (mouse, Santa Cruz).

### *In vitro* co-translation

1ug of Ajuba/Ajuba mutant plasmid and 1ug of RPA subunit/RPA70/RPA70 mutant plasmid were added into the same reaction. The *in vitro* translation protocol was performed as described in manual from manufacturer (Promega, TNT© T7 Coupled Reticuloccyte Lysate System) with 35S-methionine.

### His-tag affinity pulldown

Ni-NTA beads (BioRad, 156-0133) were washed with dH_2_O and TBST (pH=8.0, 0.01% Tween-20). The beads were blocked with 2% BSA (Sigma) (10mg/ml) in TBST for 2 hours at room temperature. 20ul of blocked beads were added to each sample and incubated at room temperature for 30 minutes. Beads were washed with 100ul TBST six times and loading buffer was added directly to beads.

### Autoradiography

SDS-PAGE gels were dried and placed for exposure into a phosphor storage cassette overnight (Amersham). The data was visualized and quantitated by Phosphor imager (Typhoon) using the ImageQuant software.

## Acknowledgments

This work was funded by a SC3 score award # 1SC3GM094071-01A1 from the National Institute of General Medical Sciences. The authors thank Danielle Khan and Baila Schochet, both Hunter College undergraduates, for excellent technical assistance, and the Inga Richter Fellowship for invaluable support to S.F.

